# Power-law Null Model for Bystander Mutations in Cancer

**DOI:** 10.1101/001651

**Authors:** Loes Olde Loohuis, Andreas Witzel, Bud Mishra

**Affiliations:** Department of Computer Science, CUNY, The Graduate Center, New York, 365 Fifth Avenue, New York, NY 10016, USA; Google, 76 Ninth Avenue, New York, NY 10011, USA; NYU Courant Institute, 251 Mercer Street, New York, NY 10012, USA

**Keywords:** Copy Number Variation, Power-law Distribution, Generative Mechanism, Cancer Driver Genes Detection

## Abstract

In this paper we study Copy Number Variation (CNV) data. The underlying process generating CNV segments is generally assumed to be memory-less, giving rise to an exponential distribution of segment lengths. In this paper, we provide evidence from cancer patient data, which suggests that *this generative model is too simplistic*, and that *segment lengths follow a power-law distribution instead*. We conjecture a simple preferential attachment generative model that provides the basis for the observed power-law distribution. We then show how an existing statistical method for detecting cancer driver genes can be improved by incorporating the power-law distribution in the null model.

## I. Introduction

COMPREHENSIVE knowledge of the genomic aberrations that underlie cancer is of vital importance for diagnostics, prognostics, and the development of targeted therapies. Towards this goal, large databases of genomic cancer-patient data are being generated in recent years. One type of such data is Copy Number Variation (CNV) data. CNV is structural variation in which relatively large regions of the genome are either amplified or deleted, leading to gain- or loss-of-function of the genes contained in the affected regions.

CNV data consists of copy-number values of thousands of markers corresponding to different locations in the genome. To reduce the noise in this data, sets of neighboring markers are often combined resulting in contiguous segments of equal copy number, classified into *normal*, *amplified*, or *deleted* segments. Examples of such tools, usually called ‘segmenters,’ include GLAD [7], CBS [11], and a method developed by Mishra’s group [2]. The abnormal segments correspond to duplication or deletion events and are used as input data to identify regions containing genes that are relevant for the development of cancer. (e.g., methods described in [9, 1]).

The underlying process generating these CNV segments is generally assumed to be memory-less, giving rise to an exponential distribution of segment lengths. In this paper, we provide evidence from cancer patient data, which suggests that *this generative model is too simplistic*, and that *segment lengths follow a power-law distribution instead*. We conjecture a simple preferential attachment generative model that provides the basis for the observed power-law distribution.

From a thorough understanding of the statistical properties of genomic copy-number data in cancer, one expects to discover (either directly or indirectly) improved oncogenomics features, using statistical inference tools which build upon more accurate null-models (examples of these tools include [7, 11, 2, 9, 1]). In this paper, we provide one such improved estimator to an existing statistical method (due to Ionita et al. [9]) for detecting genetic regions relevant to cancer, which we achieve by incorporating the power-law distribution in the null. We analyze three TCGA CNV data sets and show that the improved model based on power-law distribution outperforms the simpler null model which only uses a non-informative prior.

December 31, 2013

## II. Evidence and Fitting

We analyzed three CNV data sets from The Cancer Genome Atlas (TCGA): Lung Squamous Cell Carcinoma (LUSC 201 patients), Glioblastoma (GBM 299 patients), and Ovarian Serous Cystadenocarcinoma (OV 337 patients)^1^. The level 2 data was segmented using the segmentation algorithm of Daruwala et al. [2] and the empirical segment-length distributions of amplifications and deletions were fit to both power-law (*cx^-α^*) and exponential (*ce^-λx^*) distributions.

Figure 1 shows the segment length distribution and fitted functions for the deleted segments of the OV dataset, and Table I lists the numerical values of all fits, as well as their *R*^2^ goodness of fit. Plots for the remaining data sets can be found in figure 2 of Section A.

**Fig. 1:**
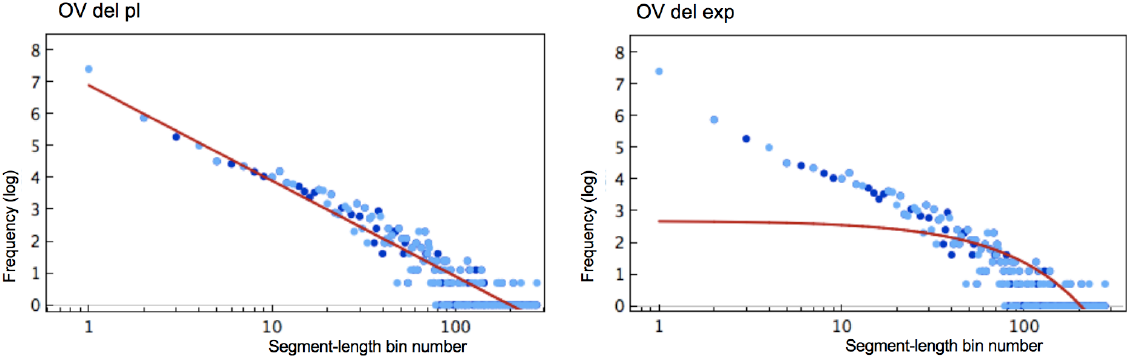
Segment length distribution and fitted functions of deleted segments from the OV dataset. The best power-law fit is shown on the left and the best exponential fit on the right. See Appendix A Figure 2 for the images showing the fits for all other data sets.

**TABLE I:** Comparison of exponential and power-law fits for three TCGA data sets: LUSC, OV, and GBM.

|  | best exponential fit |  | best power-law fit |  |
| --- | --- | --- | --- | --- |
| | function | $R^2$ | function | $R^2$ |
| LUSC Amp | $e^{-0.014x}$ | 0.65 | $x^{-1.27}$ | 0.86 |
| LUSC Del | $e^{-0.008x}$ | 0.45 | $x^{-0.89}$ | 0.79 |
| OV Amp | $e^{-0.014x}$ | 0.67 | $x^{-1.39}$ | 0.91 |
| OV Del | $e^{-0.013x}$ | 0.64 | $x^{-1.30}$ | 0.91 |
| GBM Amp | $e^{-0.015x}$ | 0.39 | $x^{-1.01}$ | 0.71 |
| GBM Del | $e^{-0.012x}$ | 0.60 | $x^{-1.20}$ | 0.78 |

To determine threshold values for amplifications and deletions, we suitably modify the method described in [8], which implies that a segment is treated as an amplification (or resp. a deletion) if its value greater (or reps. smaller) than the mean plus (or reps. minus) twice the standard distribution (*AVG ±* 2*STD*). The fit was estimated by collecting all the segment-lengths of segments above the amplification threshold value or below the deletion threshold value and taking a histogram of the segment lengths. To make the fit particularly sensitive to the tail of the distribution, we chose to fit the log of the data against the log of the exponential and power-law distributions.

As shown in Table I, in all three datasets, the power-law fits the segment-length distributions better than the exponential one.

Several remarks about this result are due at this point. First, the remaining segments that are not considered amplifications or deletions (the ‘Normals’), are not clearly power-law (nor exponentially) distributed (see Appendix A Table III for the actual fits, and Figure 3 for an illustrative figure). The power-law distribution only appears to fit segments above (or below) a certain threshold. In Appendix A, we provide some analysis of the fits relative to a selected threshold. Second, taking the logarithm of the data is a way to magnify the difference between the power-law and exponential fit, which occurs mostly in the tail. It should be noted, however, that it does not affect the relative goodness of the exponential and power-law fit, as can be verified by the results listed in Table V in the Appendix A.

### A. Generative Model

The observed power-law distributions for amplifications and deletions can be explained by a mechanism of preferential attachment. That is, once a region has large aberrations, it is more likely to acquire even more numerous large aberrations. One straightforward reason that could underlie this mechanism is that large amplifications or deletions lead to genomic instability and hence allow for subsequent large copy number aberrations.

## III. Improving tools through More Accurate Statistical Null-Models

Most of the tools that are developed to analyze genomic data assume a non-informative exponential null-model for segment length distribution (e.g., segmenters [2] and tools for detecting cancer genes [9]). Knowledge of the fact that segment lengths are not exponentially distributed allows us to improve our null models and hence our tools. This resulting prior is especially important when there is not sufficiently enough data available to accurately predict null-models from the data. In the next section we show how an existing tool for detecting cancer genes can be improved.

### A. Statistical Method for Detecting Cancer Genes

In this section we adopt a method described in [9] for finding cancer driver genes from copy number variation data by building upon the assumption that segment lengths are power-law distributed.

Cancer genes are generally divided into two types: tumor suppressor genes (TSGs) and oncogenes (OGs). TSGs prevent tumor development by regulating cell growth. A loss or reduction in its function (for example by a deletion), can lead to uncontrolled cell division and allows the cancer to progress. Oncogenes, on the other hand, are genes whose function promote proliferation. Gain-of-function mutations (like amplifications), or overexpression, promote tumor progression. In the case of TSGs a deletion of a part of the gene will cause a loss-of function, while for OGs the whole gene needs to be amplified as a whole to cause a gain-of-function.

The algorithm for finding TSGs and OGs enumerates all possible intervals and assigns to them a score function that measures the likelihood of this being a driver gene. This score function can be described as follows:

For any interval *I* the strength of the association between deletions in *I* or amplifications of *I* and the disease is quantified by analyzing the genomic data for many individuals with a specific type of cancer. For this purpose, a metric called *Relative Risk* (*RR* _event_ *_I_*) assigns a numerical value to any event, a deletion or amplification of an interval, which thus compares the probability of the disease occurring with or without the event. Informally, *RR* _event_ *_I_* is the degree to which the occurrence of event *I raises the probability* of the disease incidence. Formally,

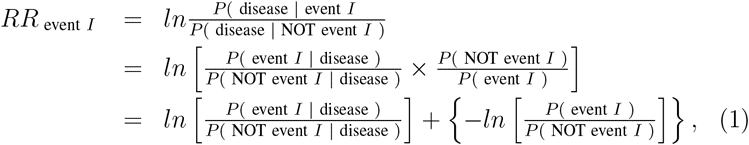

where, in case of a deletion, “event *I*” denotes the event that *at least part* of *I* is deleted. We call this event ‘*I* broken’. In case of an amplification “event *I*” denotes the event that there exists an amplified interval that *fully includes I*. We call this event ‘*I* increased’.

The first term in equation (1) can be computed from the available tumor samples:

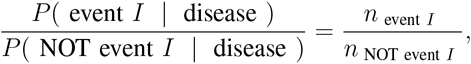

where *n* _event_ *_I_* (or *n* _NOT event_ *_I_*) is the number of patients in whose tumor genomes the event *I* occurs (or does not occur). Note that becasue of the intrinsic differences between TSGs and OGs in case of deletions, the longer the segment the larger 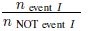 whereas in case of amplifications the situation is reversed: longer segments have smaller 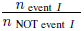. This imbalance is corrected for by the second part of (1),

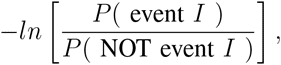

which incorporates prior information inherent in the statistical distribution of amplifications and deletions.

To compute the prior score, we assume that, at any genomic location, a breakpoint (starting point) may occur as a Poisson process at a rate of *µ ≥* 0. We consider two different *µ*’s: one for amplifications *µ*_AMP_ and the other for deletions *µ*_DEL_, but we drop the subscript when no confusion arises. Segments are modeled as vectors. Starting at a breakpoint and moving left (or right) with probability 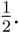 The length *t* of each segment is distributed according to a power-law distribution: *t^-α^*, with 1 *≤ α ≤* 2. Let *ϵ* be the constant that represents the shortest length an interval could possibly have.

Given these assumptions we can derive the prior probability that an interval *I* is amplified or deleted.

#### Proposition III.1.

*Assuming that segment lengths are power-law distributed:*

1. *The probability that an interval I* = [a, b] *is broken is as follows:*

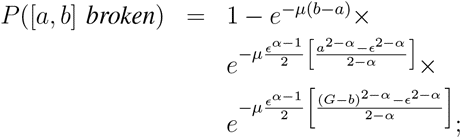
2. *The probability that an interval I* = [a, b] *is increased is as follows:*

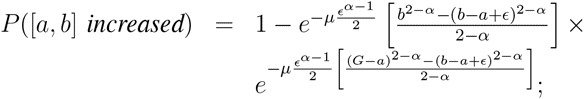

*where* [0*, G*] *represents the region of interest (e.g. a chromosome) and* [*a, b*] *is an interval within this region. It is assumed that ϵ ≪ G*.

The proof of this proposition can be found in Appendix III.1.

The parameter *α* can be estimated from the data as described in section II. The values of the *µ*_DEL_ and *µ*_AMP_ parameters are the mean number of amplifications and deletions per unit length respectively and can be computed directly from the segmented data.

The constant *ϵ* can take any value. If we assume the value of *ϵ* is 1 unit (corresponding to a single probe in microarray data or a single base in sequencing data) the probability that a segment is broken approaches:

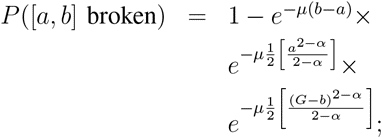

Similarly for amplifications:

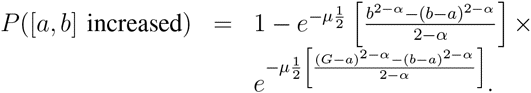

The *RR* score can be used to estimate the location of tumor suppressor genes and oncogenes. The simplest algorithm first computes the score for all intervals with value in a range determined by lower and upper bounds, and then picks the highest scoring interval on each chromosome. Many other algorithms can be imagined. For example, one can use two scoring functions to compute the left and right boundaries of the interval separately. The final step of the algorithm is significance testing of the obtained intervals. The methods as described in [9] for tumor suppressor genes, and in [8] for oncogenes can be directly applied. Both methods assign a *p*-value for every putative TSG or oncogene using tools from *scan statistics* [12].

We have implemented the algorithm by computing the RR score for each interval while keeping track of the highest scoring interval. Because each interval needs to be visited only once the time complexity is linear in the number of intervals.

Instead of finding only the interval with maximum score on each chromosome we can let the algorithm pick higher scoring intervals. One straightforward way is to pick the *n* non-overlapping significantly amplified/deleted intervals with the highest score, by keeping track of a list of results while going through the set of all intervals. This method has certain shortcomings as described in the discussion section.

### B. Performance Comparison

To be able to test the influence of the improved null model, we have applied the afore-described algorithm with both the original exponential and the power-law null models to the three TCGA datasets: OV, LUSC and GMB.

To compare the two models we asked which of the commonly amplified or deleted genes in the three cancer types were found by the respective algorithms. The results are summarized in table II Consistent with our expectation, the power-law based model performs (slightly) better than the exponential model.

**TABLE II:** List of genes that are commonly altered in OV, LUSC and GBM cancer cells, and whether or not they were found by the power-law and exponential methods using the three highest scoring non-overlapping intervals. A more detailed version of this table can be found in Table VIII in Appendix C.

| Cancer | Gene | Power-law | Exponential |
| --- | --- | --- | --- |
| OV | BRCA1 | no | no |
|  | BRCA2 | no | no |
|  | ERBB2 | no | no |
|  | K-ras | **yes** | **yes** |
|  | AKT2 | no | no |
|  | PIK3CA | no | no |
|  | c-MYC | **next** | **no** |
|  | p53 | no | no |
| LUSC | CDKN2A | **yes** | **yes** |
|  | FGFR1 | **yes** | **no** |
|  | PDGFRA | no | no |
|  | SOX2 | no | no |
|  | HWSCL1 | **next** | **no** |
| GBM | EGFR | **next** | **next** |
|  | MDM2 | no | no |
|  | PDGFR | no | no |
|  | CDK4 | no | no |
|  | Rb | no | no |
|  | CDKN2A | **yes** | **yes** |

Note that despite the (slightly) better performance of the algorithm with the power-law null model over the exponential model, the difference between the two performances is comparable and both algorithms appear to miss many cancer genes. Both methods can be further improved by including additional information (e.g., gene-ontologies, gene-networks or pathways). In such a setting, as well as when regions for many more genes are checked, the contribution from more accurate null model is expected to be more pronounced.

We offer several explanations for the missing genes. For example, the algorithm only picks out a few (in this case three) high scoring intervals per chromosome. Often, these intervals are in the same region close to a single gene, which causes other regions of interest to be overlooked. For example, in the OV dataset, all three deleted intervals that were found on chromosome 17 were close to (but not exactly overlapping with) BRCA1. It became therefore impossible to find P53, which also lies on chromosome 17, as well. This problem can be resolved by adopting more sophisticated statistical methods for selecting high-scoring intervals.

In addition, regions either right next to actual genes or close to the centromere were often identified as likely cancer genes. We expect this type of error to disappear as methods for CNV data collection become more precise. In the next section, we briefly mention several other possible ways to improve the method for finding driver genes.

## IV. Conclusions and Discussion

In summary, we have provided evidence suggesting that the segment lengths of CNV amplifications and deletions in cancer cells follow a power-law distribution instead of the commonly assumed exponential distribution. This evidence suggests a generative mechanism of preferential attachment: many long amplifications and deletions lead to even more long amplifications and deletions. Even though our data analysis rules out exponentially distributed segment lengths, and the evidence for power-law distribution is compelling, other distributions (such as log-normal or stretched exponential, see Table VI in Appendix A) cannot be completely excluded on the basis of this evidence.

Especially in cases where only a small sample of data is available to estimate the prior distribution from the data, knowledge about the statistics of CNV data allows us to improve our analytic tools. As an example, we have demonstrated how the technique for finding cancer driver genes described in [9] can be modified to incorporate the power-law distribution and, as our preliminary results indicate, how the power-law-based scan-statistics algorithm outperforms the exponential one. Once inferred, the set of cancer driver genes can be used as input to cancer progression extraction algorithms to derive progression models from static cancer patient data (see e.g., [3, 4, 6, 5]), leading to improved diagnostics, prognostics, and targeted therapies.

We note in conclusion that, despite its promise, these results represent an analysis that remains largely preliminary in nature. More recent single-cell single molecule genomic data have shed light on the significant heterogeneity and temporality that exist in cancer progression – namely, a tumor consists of a heterogeneous population of cell-types and the cells of different cell-types interact dynamically going through rapidly-changing cell-states. Thus, more sophisticated oncogenomic analysis tools will need to generalize the mathematics described here much further, in which the null model must include a mixture of distributions, with the parameters of the distribution fluctuating as cancer progresses. Consequently, the tool to find cancer driver genes can be further improved in several ways. For example, we will need to incorporate a preferential attachment model to the segmenter that analyzes the genomic data from each cell-type; use more accurate priors of the *distribution of breakpoints* that are known to occur in different cell-types; apply more sophisticated statistical tools for picking high-scoring intervals by incorporating prior biological knowledge (carefully, so as to avoid Bayesian bias); and include such information (i.e., how pathways affect the cell-states) in combination with precise correction for multiple hypothesis testing in order to make the final results more meaningful. But, to keep the focus on just the algorithmic/mathematical nature of this problem, the formulation developed here has been kept rudimentary; thus, a more practical description of a complete solution has remained outside the scope of this paper.

## Appendix A Segment-length distribution

Let *AVG_C_* and *STD_C_* (resp *AVG_N_* and *STD_N_*) denote the average segment-length and the standard deviation of all segments derived from tumor (resp blood-derived normal) cells.

**Fig. 2:**
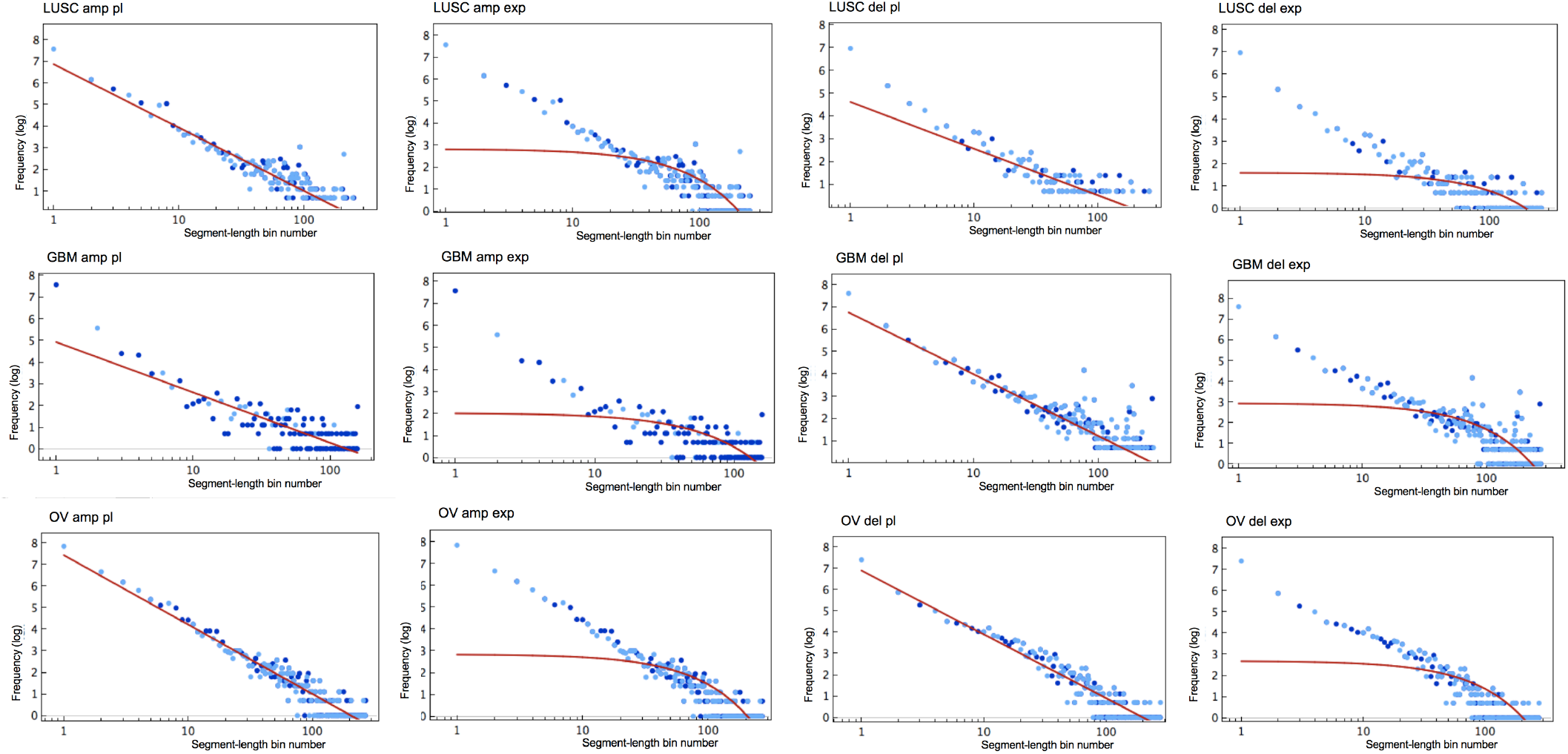
Segment length distribution and fitted functions for all three datasets: LUSC, OV, GBM. The thresholds are *AVG_C_ ±* 2*STD_C_*.

**Fig. 3:**
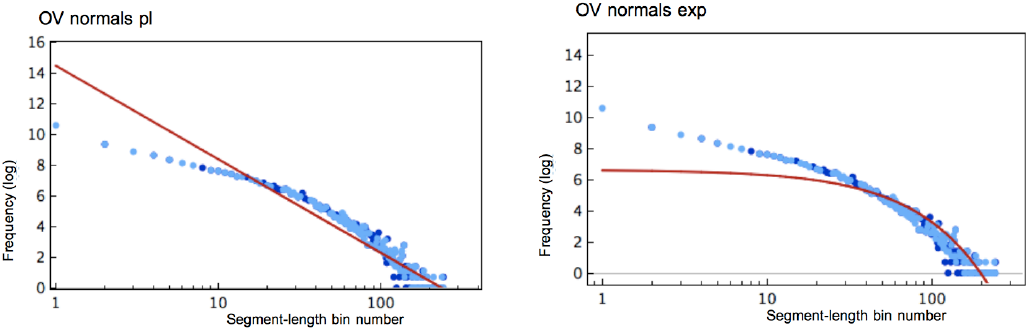
Segment length distribution and fitted functions for OV ‘Normals’. That is, all segments with segment values in [*AVG_C_ −* 2*STD_C_, AVG_C_* + 2*STD_C_*].

**TABLE III:**
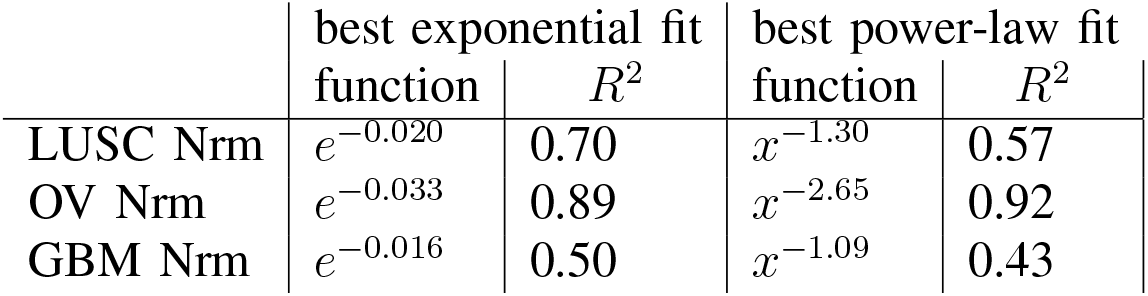
Distribution fits of the ‘Normals’.

**TABLE IV:** Using the OV dataset, this table shows how different tresholds influence the power-law and exponential fits.

| Threshold | OV |  |  |  |  |  |  |  |  |  |
| --- | --- | --- | --- | --- | --- | --- | --- | --- | --- | --- |
|  | AMP |  |  |  |  | DEL |  |  |  |  |
|  | th | PL |  | EXP |  | th | PL |  | EXP |  |
| | | $\alpha$ | $R^2$ | $\lambda$ | $R^2$ | | $\alpha$ | $R^2$ | $\lambda$ | $R^2$ |
| $\pm 1.0$ | 1.00 | 1.41 | 0.90 | 0.024 | 0.64 | -1.00 | 1.20 | 0.85 | 0.015 | 0.52 |
| $AVG_C \pm 2STD_C$ | 0.76 | 1.39 | 0.91 | 0.014 | 0.67 | -0.87 | 1.30 | 0.91 | 0.013 | 0.64 |
| $AVG_C \pm 1.5STD_C$ | 0.59 | 1.79 | 0.93 | 0.031 | 0.81 | -0.66 | 1.82 | 0.93 | 0.028 | 0.75 |
| $AVG_C \pm 1STD_C$ | 0.36 | 1.94 | 0.93 | 0.030 | 0.84 | -0.46 | 1.99 | 0.90 | 0.033 | 0.85 |
| $AVG_N \pm 5STD_N$ | 0.21 | 2.11 | 0.88 | 0.0251 | 0.85 | -0.27 | 2.21 | 0.85 | 0.0343 | 0.87 |
| $AVG_N \pm 3STD_N$ | 0.11 | 1.85 | 0.85 | 0.0255 | 0.87 | -0.18 | 1.99 | 0.86 | 0.0343 | 0.89 |
| $AVG_N \pm 2STD_N$ | 0.06 | 1.79 | 0.83 | 0.025 | 0.89 | -0.13 | 1.96 | 0.86 | 0.034 | 0.90 |
| 0.0 | 0.00 | 1.79 | 0.83 | 0.025 | 0.89 | 0.00 | 1.96 | 0.86 | 0.034 | 0.90 |

**Fig. 4:**
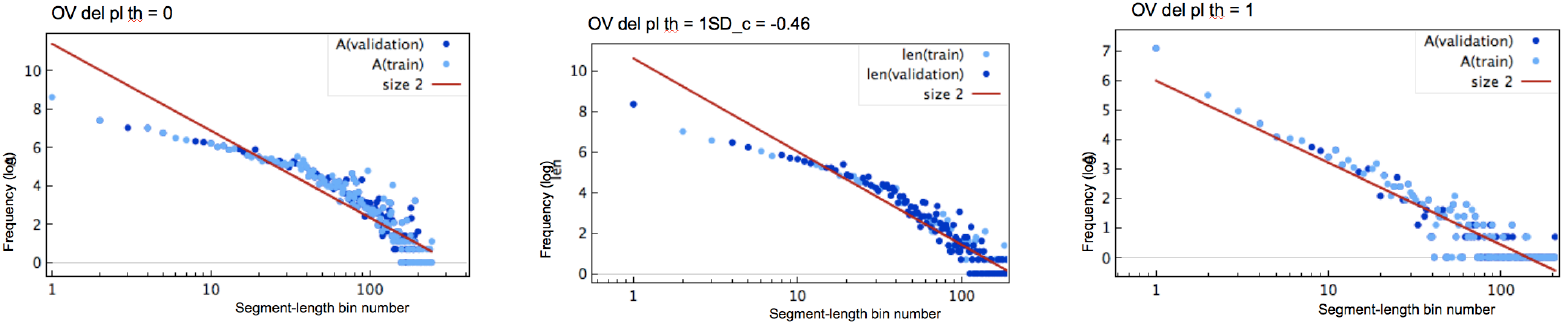
OV deletions segment length distributions for different thresholds: 0, *AVG_C_* ± 1 *SD_C_* = −0.46 and −1.

**Fig. 5:**
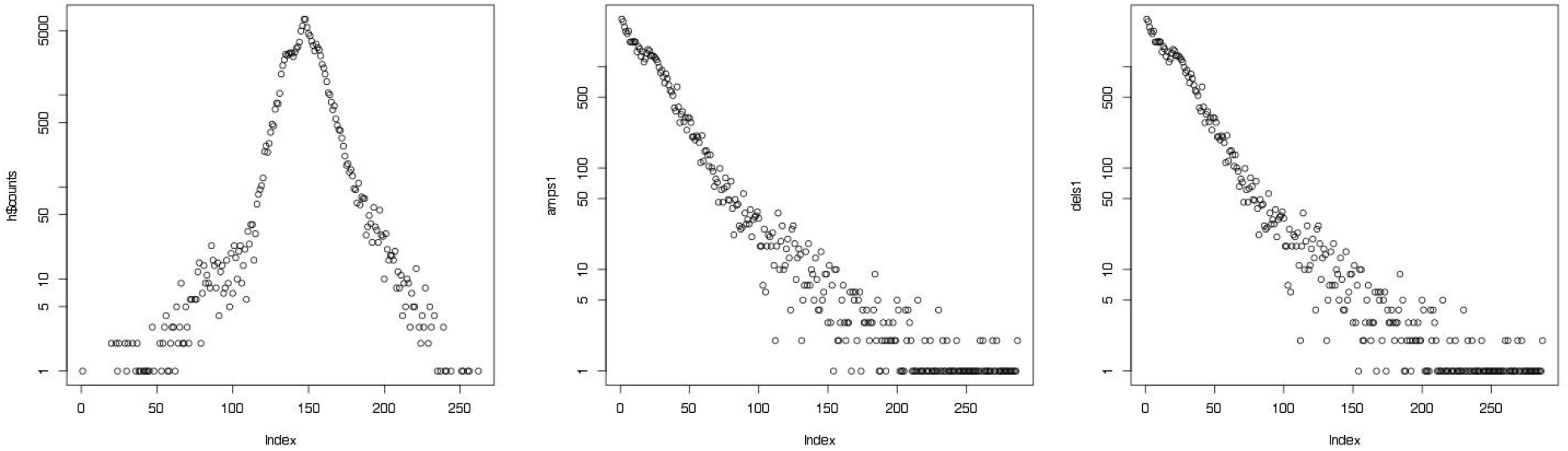
Distribution of segment values of all segments (left), all positive segment values (middle) and negative segment values (right), on a log-log scale.

## Appendix B Proof of proposition III.1

**The Model:** We assume that, at any genomic location, a breakpoint (starting point) may occur as a Poisson process at a rate of *µ ≥* 0. We consider two different *µ*’s: one for amplifications *µ*_AMP_ and one for deletions *µ*_DEL_, but we drop the subscript when no confusion arises. Segments are modeled as vectors. Starting at a breakpoint *x* and moving left (or right) with probability 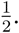 The length *t* of each segment is distributed according to a power-law distribution: *t^-α^*, with 1 *≤ α ≤* 2. Let *ϵ* be the constant that represents the shortest length an interval could possibly have.

**TABLE V:** Exponential and power-law fits for non-log data.

| | best exponential fit<br>function | $R^2$ | best power-law fit<br>function | $R^2$ |
| --- | --- | --- | --- | --- |
| LUSC Amp | $e^{-0.27x}$ | 0.96 | $x^{-1.26}$ | 0.97 |
| LUSC Del | $e^{-0.48x}$ | 0.94 | $x^{-1.58}$ | 0.99 |
| OV Amp | $e^{-0.26x}$ | 0.96 | $x^{-1.50}$ | 0.99 |
| OV Del | $e^{-0.30x}$ | 0.91 | $x^{-1.22}$ | 0.99 |
| GBM Amp | $e^{-1.51x}$ | 1.00 | $x^{-2.71}$ | 1.00 |
| GBM Del | $e^{-0.41x}$ | 0.91 | $x^{-1.39}$ | 0.97 |

**TABLE VI:** Exponential (*ce^-λx^*), power-law (*cx^-α^*) and log-normal 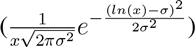 fits.

| | best exponential fit<br>$R^2$ | best power-law fit<br>$R^2$ | best log-normal fit<br>$R^2$ |
| --- | --- | --- | --- |
| LUSC Amp | 0.65 | 0.86 | 0.82 |
| LUSC Del | 0.45 | 0.79 | 0.77 |
| OV Amp | 0.67 | 0.91 | 0.77 |
| OV Del | 0.64 | 0.91 | 0.86 |
| GBM Amp | 0.39 | 0.71 | 0.71 |
| GBM Del | 0.60 | 0.78 | 0.76 |

### Proposition B.1.

*Assuming that segment lengths are power-law distributed:*

1. *The probability that an interval I* = [*a, b*] *is broken is as follows:*

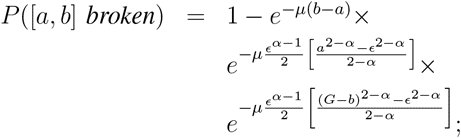
2. *The probability that an interval I* = [*a, b*] *is increased is as follows:*

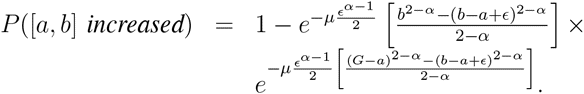 *Proof:*

1. We wish to estimate the probability that the interval [*a, b*] is ‘broken’. This is the probability that there exists a deleted interval *I* that intersects with [*a, b*]:

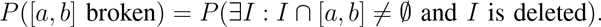 Instead, we compute *P*([*a, b*] is NOT broken) by computing:

(*P*_1_) The probability that no deletion occurs starting in the interval [*a, b*],
(*P*_2_) The probability that each deletion starting in [0*, a*] does not overlap [*a, b*], and
(*P*_3_) The probability that each deletion starting in [*b, G*] does not overlap [*a, b*]. It follows that *P* ([*a, b*] is NOT broken) = *P*_1_ *× P*_2_ *× P*_3_. Thus,

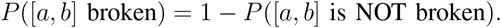

(*P*_1_) *P*(no deletion starts in [*a, b*]) = *e^-µ^*^(*b-a*)^. This equation follows immediately from the assumption that breakpoints are generated by a Poisson process. Note that we drop the subscript *DEL* in *µ*_DEL_.
(*P*_2_) *P*(each interval starting in [0*, a*] does not overlap with [*a, b*]) can be broken down as the following infinite sum: *P* (each interval starting in [0*, a*] does not overlap with [*a, b*]) =

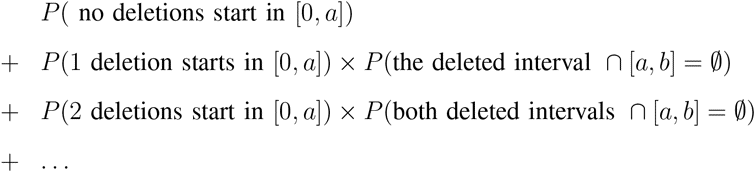 By the assumption that breakpoints are generated as a Poisson process, the probability *P*(*n* deletions start in 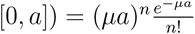 for each *n*. The probability *P*(1 deleted interval∩[*a, b*] = *∅*) can be computed as follows. From our model it follows that *P*(deleted interval *∩* [*a, b*] = *∅ |* 1 deletion starts in [0*, a*]) is the probability that each deletion starting at *x* in the interval [0*, a*] does not reach all the way to *a*: *P* (deleted interval *∩* [*a, b*] = *∅ |* 1 deletion starts in [0, *a*])

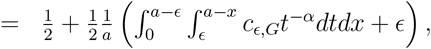

where:

– The constant *c*_*ϵ*, *G*_ depends on the length *ϵ* and *G* and is computed below.
– The 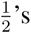 are to take into account the possibility that the deletion moves left instead of right.
– The last term + *ϵ* takes into account the possibility that the starting point of the deleted interval is in [*a – ϵ, a*]. The preceding equation can be simplified as follows:

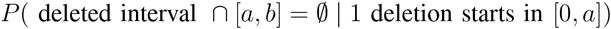

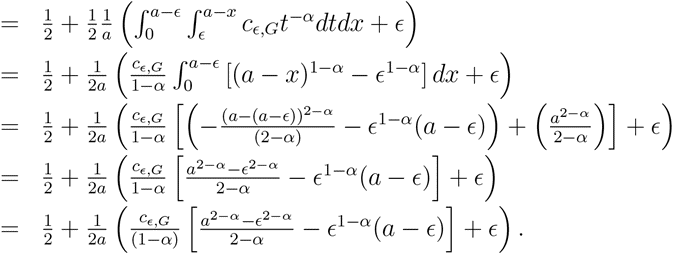 There are a few points to make regarding this derivation:

– We ignore the integration constants, as they cancel each other out.
– Since *α ≥* 1, and *c*_ϵ_, _*G*_, *a ≥* 0, the term 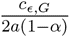 is negative. We thus need to show that 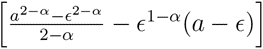 is always negative to obtain a positive probability. This follows from the mean-value theorem. Namely, for any function *f* that is concave and increasing the following holds:

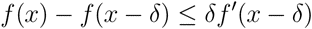

the function 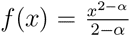 is concave and increasing with *f ^′^*(*x*) = *x*^1*-α*^. If we let *x* = *a* and *δ* = *a – ϵ* then *x – δ* = *ϵ* and we have

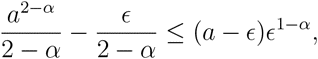

from which it follows that 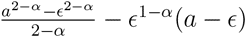 is negative. The normalizing constant *c_ϵ_* *_,G_* can be computed as follows. It has to be such that

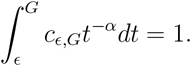 It follows that

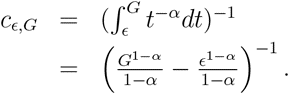 Since *α >* 1 and *G ≫ ϵ* this approaches

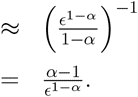 Using 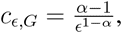 we can simplify *P* (deleted interval *∩* [*a, b*] = *∅ |* 1 deletion starts in [0*, a*]) as follows:

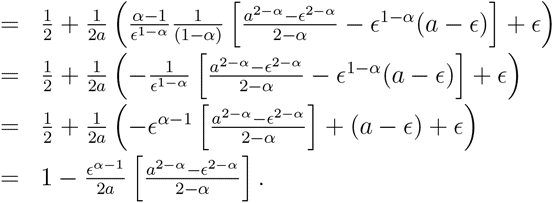 Since deletions are assumed to be independent events that can overlap it follows that *P* (*n* deleted intervals *∩* [*a, b*] = *∅ | n* deletions starts in [0*, a*]) = *P* (deleted interval *∩*[*a, b*] = ∅ | 1 deletion starts in [0*, a*])*^n^* Hence, we get the following series:

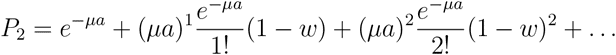

with 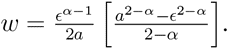 It follows that

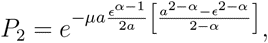

which can be simplified to

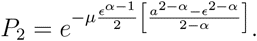 (*P*_3_) *P* (each interval starting in [*b, G*] does not overlap with [*a, b*]) is computed in the same way as *P*_2_, but now starting at *x* ∈ [*b, G*] and moving left. In this case

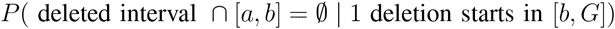

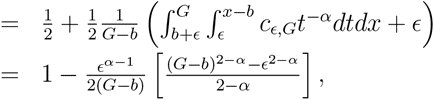

and we obtain

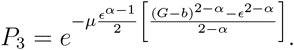 It follows that

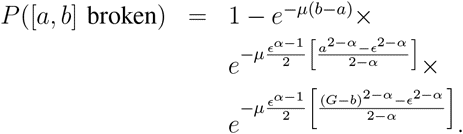
(2) In an analogous fashion, we calculate the probability that the interval [*a, b*] is ‘increased’. This is the probability that there exists a deleted interval *I* that includes [*a, b*]:

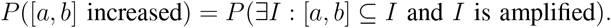 We compute *P* ([*a, b*] is NOT increased) by computing:

(*P*_1_) The probability that each interval starting in [0*, a*] does not include [*a, b*], and
(*P*_2_) The probability that interval starting in [*b, G*] does not include [*a, b*]. The computation of *P*_1_ (and *P*_2_) is exactly like that of deletions, except for the fact that we can to integrate over all intervals reaching up to *b* (down to *a*). In the case of *P*_1_, we solve 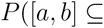 amplified interval | 1 amplification starts in [0*, a*])

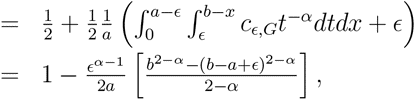

and in the case of *P*_2_ 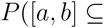 amplified interval | 1 amplification starts in [*b, G*])

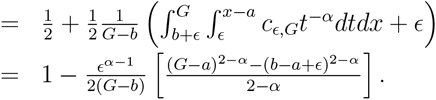 We obtain:

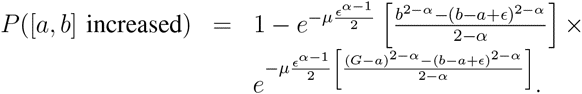

## Appendix C Detecting Driver Genes

**TABLE VII:**
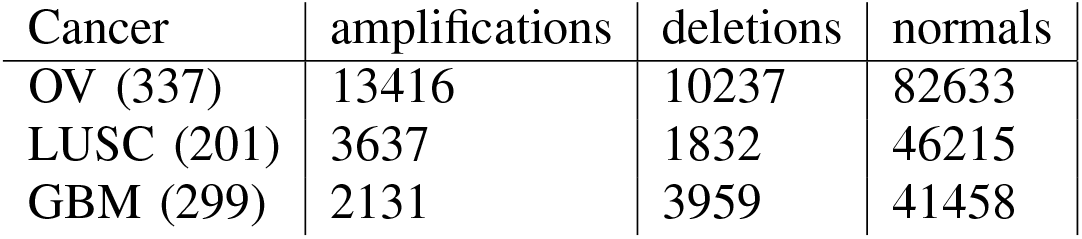
Number of deleted and amplified segment for three TCGA data sets using a threshold of *AVG_C_ ±* 2*STD_C_*.

**TABLE VIII:** List of genes with their locations that are commonly altered in OV, LUSC and GBM cancer cells, and whether or not they were found by the power-law and exponential methods using the three highest scoring non-overlapping intervals. The OV and GBM genes were taken from the Kegg database (http://www.genome.jp/dbget-bin/www_bget?ds:H00027 and http://www.genome.jp/dbget-bin/www_bget?ds:H00042); the LUSC genes, for which no Kegg entry exists, are commonly amplified/deleted LUSC driver genes from [10] (mentioned on page 519). A gene is considered ‘found’ if the selected interval intersects with the region containing the gene. In this table ‘next’ indicates within 100kbp from a border of the gene interval. The parameters *µ*, *α*, and *λ* were estimated from the data as in [9] and Table I, with the exception of *α* of LUSC del which was set to 1, as the computation of RR score assumes *α ≥* 1. Segments shorter than 10^4^ base pairs (corresponding to the distance between two probes) and longer than 10^7^ base pairs were excluded.

| Cancer | Gene | Function | Location | Power-law | Exponential |
| --- | --- | --- | --- | --- | --- |
| OV | BRCA1 | TSG | 17: (41196312..41277500) | no | no |
|  | BRCA2 | TSG | 13: (32889617..32973809) | no | no |
|  | ERBB2 | OG | 17: (37844393..37884915) | no | no |
|  | K-ras | OG | 12: (25358180..25403854) | yes (25289555..25421243) | yes (25177510..26726740) |
|  | AKT2 | OG | 19: (40736224..40791302) | no | no |
|  | PIK3CA | OG | 3: (178866311..178952500) | no | no |
|  | c-MYC | OG | 8: (128748315..128753680) | next (128797789..128989029) | no |
|  | p53 | TSG | 17: (7571720..7590868) | no | no |
| LUSC | CDKN2A | TSG | 9: (21967751..21994490) | yes (18947155..28723296) | yes (21983401..21993651) |
|  | FGFR1 | OG | 8: (38268656..38326352) | yes (38303346..38369274) | no |
|  | PDGFR | OG | 4: (55095264..55164412) | no | no |
|  | SOX2 | OG | 3: (34650005..34652461) | no | no |
|  | WHSC1L1 | OG | 8: (38132560..38239790) | next (38303346..38369274) | no |
| GBM | EGFR | ONCG | 7: (55086725..55275031) | next (55049021..55065490) | next (54998411..55043660) |
|  | MDM2 | ONCG | 12: (69201971..69239320) | no | no |
|  | PDGFR | ONCG | 5: (149493402..149535422) | no | no |
|  | CDK4 | ONCG | 12: (58141510..58146230) | no | no |
|  | Rb | TSG | 13: (48877883..49056026) | no | no |
|  | p53 | TSG | 17: (7571720..7590868) | no | no |
|  | PTEN | TSG | 10: (89623195..89728532) | no | no |
|  | CDKN2A | TSG | 9: (21967751..21994490) | yes (21973069..21983401) | yes (21973069..21983401) |

## Footnotes

1 http://cancergenome.nih.gov/ The datasets used are: LUSC HMS_HG-CGH-415 K_G4124A, GBM HMS_HG-CGH-244A, and OV HMS_HG-CGH-415K_G4124A.

